# What is the Best Available Science?: Conservation Status of Two California Desert Vertebrates

**DOI:** 10.1101/342048

**Authors:** Adam G. Clause, Christopher J. Norment, Laura Cunningham, Kevin Emmerich, Nicholas G. Buckmaster, Erin Nordin, Robert W. Hansen

## Abstract

Scientific progress depends on evidence-based research, and reliance on accurate scholarship is essential when making management decisions for imperiled species. However, erroneous claims are sometimes perpetuated in the scientific and technical literature, which can complicate policy and regulatory judgments. The literature associated with two enigmatic California desert vertebrates, the Panamint alligator lizard *Elgaria panamintina* and the Inyo Mountains salamander *Batrachoseps campi*, exemplifies this problem. We produced a comprehensive threat analysis and status assessment for these species, which are both under review for possible listing under the US Endangered Species Act (ESA). Despite uncertainties and limited data, we find that many sources contain factual errors about the status of these two species, particularly the original petition that advocated for ESA listing. Although localized declines may have gone undetected, no evidence exists of population declines, population extirpation, or population-scale habitat conversion for *E. panamintina*. However, there is evidence of recent flash flood damage to some occupied *B. campi* habitat, which has possibly led to population declines at those localities. Contrary to inaccurate statements by some authors, all known populations of both species occur exclusively on federal lands, and numerous populations have likely benefited from recent federal management targeted at reducing known threats. Of the 12 threats that we identified for one or both species, only three currently appear to be serious: water diversions, climate change, and flash floods. The remaining threats are neither widespread nor severe, despite numerous contrary yet poorly supported statements in the literature. We thus evaluate the contemporary conservation status of both species as relatively secure, although *B. campi* is more at-risk compared to *E. panamintina*. This conclusion is independently supported by a recent review. Nonetheless, ongoing stewardship of these species in a multi-use context by federal agencies remains vital, and we identify several priority management actions and research needs for both species. We also recommend updated determinations on the IUCN Red List, and the Species of Conservation Concern list of the Inyo National Forest. To maximize the quality and effectiveness of conservation planning, we urge government agencies, non-governmental organizations, and individual scientists to maintain high standards of scholarship and decision-making.

## Introduction

Science is an incremental, evidence-based process whereby new research builds on earlier work. A common metaphor for this process is that individual works are like bricks, which are progressively assembled into structures that represent bodies of theory and descriptive knowledge (Forscher 1963; Courchamp and Bradshaw 2017). Building good “bricks” and “structures” depends on proper interpretation of prior research, comprehensive review of relevant sources, synthetic data analysis, and placement of new findings in appropriate context. Failure to adhere to these scholarly principles can misdirect scientific progress. Such issues have motivated several recent reminders of author best-practices (Perry 2016; Anonymous 2017), and the life sciences have not escaped these problems (Grieneisen and Zhang 2012). For example, narratives on the impact of invasive species are sometimes affected by inaccurate or misinterpreted citations (Stromberg et al. 2009; Ricciardi and Ryan 2017), perspectives on predator-mediated trophic cascades may be overly simplistic and even misleading (Kauffman et al. 2010; Marshall et al. 2013; Marris 2014), and poorly documented claims of both species rediscovery and species extinction/extirpation are regularly falsified (Ladle et al. 2009; Roberts et al. 2010; Ladle et al. 2011; Scheffers et al. 2011; Caviedes-Solis et al. 2015; Clause et al. 2018).

Reliance on quality scholarship is especially important when making conservation decisions for imperiled species. It ensures the best possible justification for the decision (Sutherland et al. 2004), and can increase the legitimacy of the decision among stakeholders (Pullin and Knight 2009). This precept is codified in the decision-making process associated with what is, arguably, the most far-reaching piece of environmental legislation in the United States: the US Endangered Species Act (ESA 1973, as amended). Administered by the United States Fish and Wildlife Service (USFWS) and the National Marine Fisheries Service, the ESA requires both agencies to consider only the “best scientific and commercial data available” when making species-listing decisions (ESA 1973). However, the degree to which these agencies follow this standard is variable, due to numerous internal and external challenges (Lowell and Kelly 2016; Murphy and Weiland 2016). Problems attributed to poorly supported ESA listing petitions have also motivated recent regulatory changes to the process for proposing new additions to the Lists of Endangered and Threatened Wildlife and Plants (USFWS et al. 2016).

In 2012, the US nonprofit Center for Biological Diversity submitted a multi-species petition to the USFWS advocating ESA listing for 53 amphibian and reptile taxa (Adkins Giese et al. 2012). As required under the ESA, the USFWS subsequently released 90-day findings for all 53 taxa, which represented the agency’s initial decision on whether the petitioner offered substantial information in support of listing (USFWS 2015a, 2015b, 2015c; USFWS 2016a, 2016b, 2016c). These 90-day findings concluded that, for 17 of the 53 species, Adkins Giese et al. (2012) did not present substantial information that the petitioned action (ESA listing) was warranted. The remaining 36 taxa were advanced to the status review phase for more detailed examination and public comment. Two of the taxa currently undergoing status review are the Panamint alligator lizard *Elgaria panamintina*, and the Inyo Mountains salamander *Batrachoseps campi* (Figure 1; USFWS 2015c).

**Figure 1.**
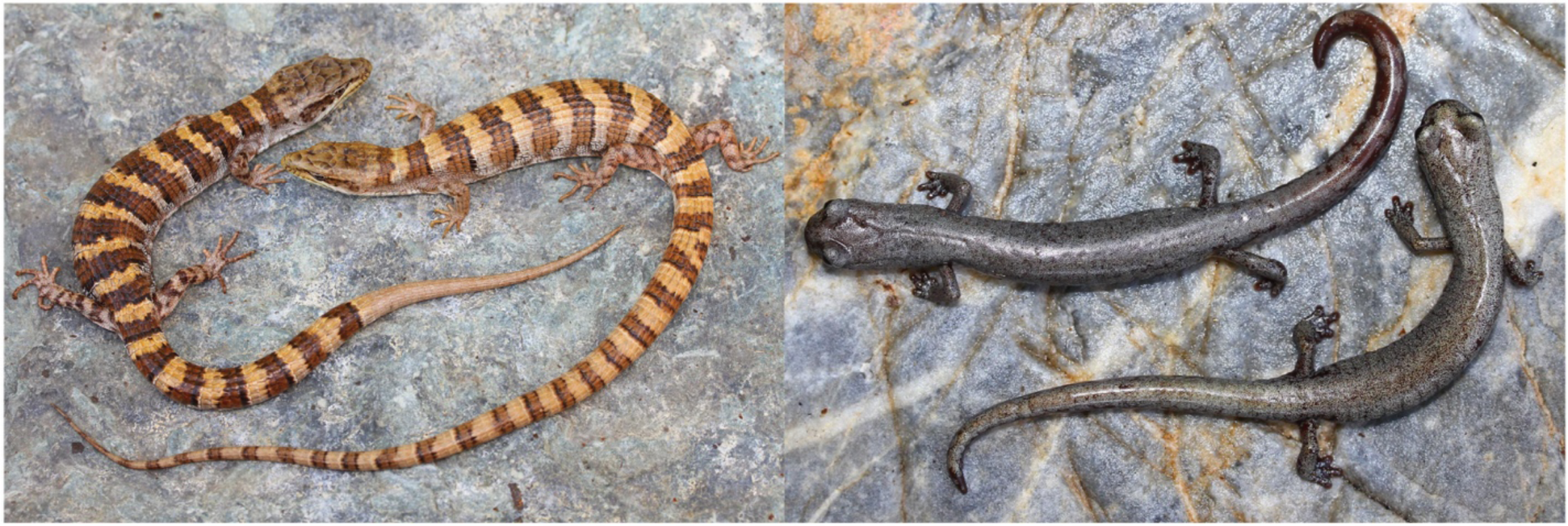
Panamint alligator lizard, *Elgaria panamintina* (left), and Inyo Mountains Salamander, *Batrachoseps campi* (right). Photographs by Adam G. Clause.

These two species are endemic to eastern California, USA, where they are roughly codistributed in the arid mountain ranges of the western Great Basin and northern Mojave deserts (Banta et al. 1996; Jockusch 2001). These mountains are among the most rugged and inaccessible landscapes in California. They support few paved roads, and their slopes are often incised by steep canyons with multiple waterfalls (Figure 2, Figure 3). Within the mountains, *E. panamintina* and *B. campi* inhabit similar environs and sometimes occur in syntopy. Occupied microhabitats for both species include mesic riparian zones fed by perennial springs or creeks, and more arid talus slopes or limestone rock crevices far from standing water (Macey and Papenfuss 1991a, 1991b). Due to their secretive behavior and remote habitats, little is known of these species’ biology and minimal literature has accrued since their discovery in 1954 and 1973, respectively (Stebbins 1958; Marlow et al. 1979). Nonetheless, both species are widely considered imperiled to some degree. The IUCN Red List of Threatened Species categorizes *E. panamintina* as Vulnerable (Hammerson 2007) and *B. campi* as Endangered (Hammerson 2004a). They have been designated as Species of Special Concern by the California Department of Fish and Wildlife (CDFW) for over 20 years (Jennings and Hayes 1994), and retained that status following a recent review (Thomson et al. 2016). However, some information that was incorporated into these determinations is inaccurate. Furthermore, many erroneous claims exist in the literature for both species, and substantial field survey data have accumulated since these listings were released. Identifying these errors, and accounting for new data, are especially important given the major regulatory decision that is pending for both species.

**Figure 2.**
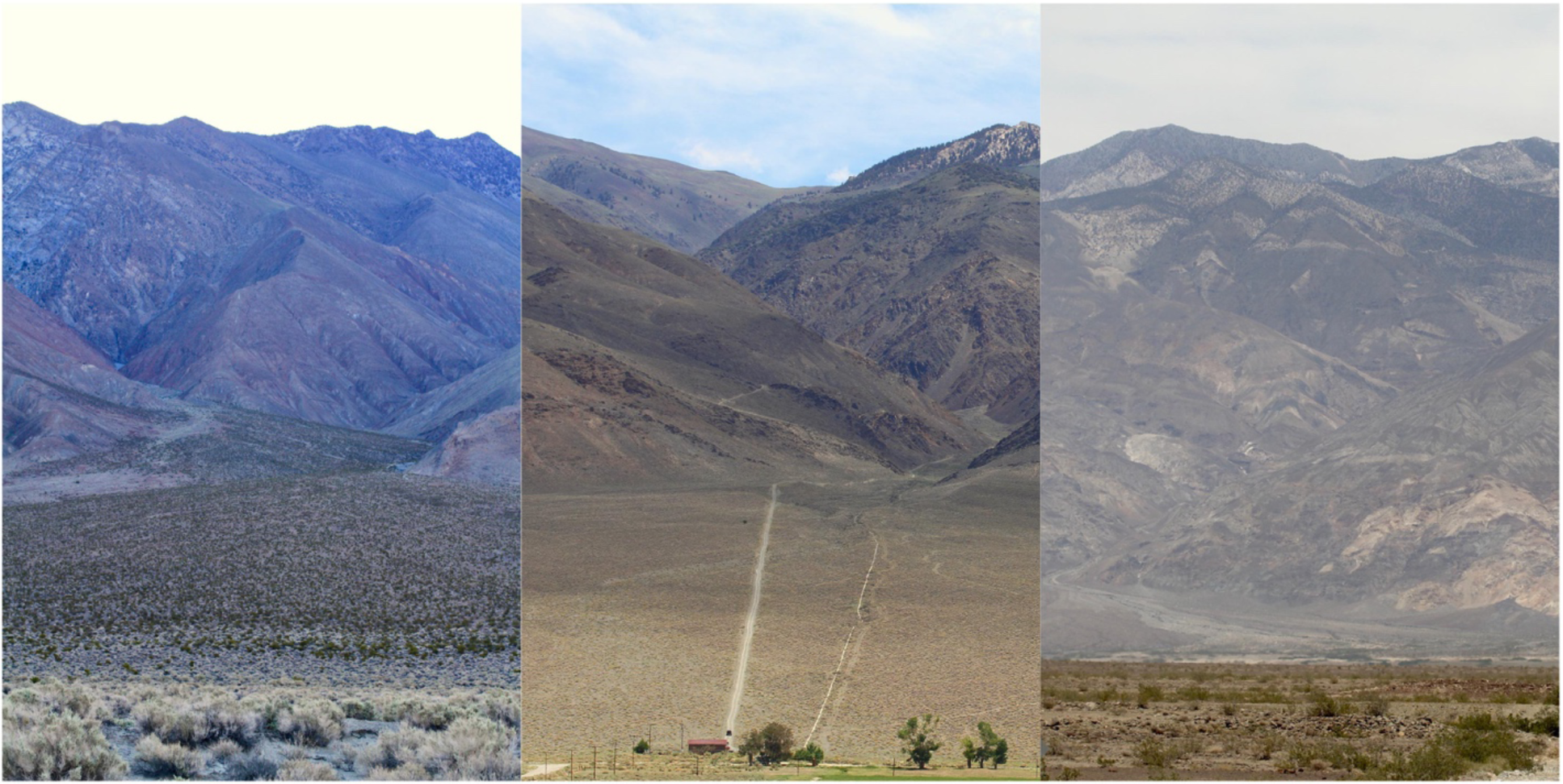
Representative landscapes for *Elgaria panamintina* and/or *Batrachoseps campi* showing the undeveloped character of the mountainous regions they inhabit. Left to right: Union Wash, Inyo Mountains; Piute Creek, White Mountains; Surprise Canyon, Panamint Mountains. Photos taken from the approximate vantage point of the nearest paved road; a high-clearance dirt access road is visible in the middle photo. Photographs by Adam G. Clause.

**Figure 3.**
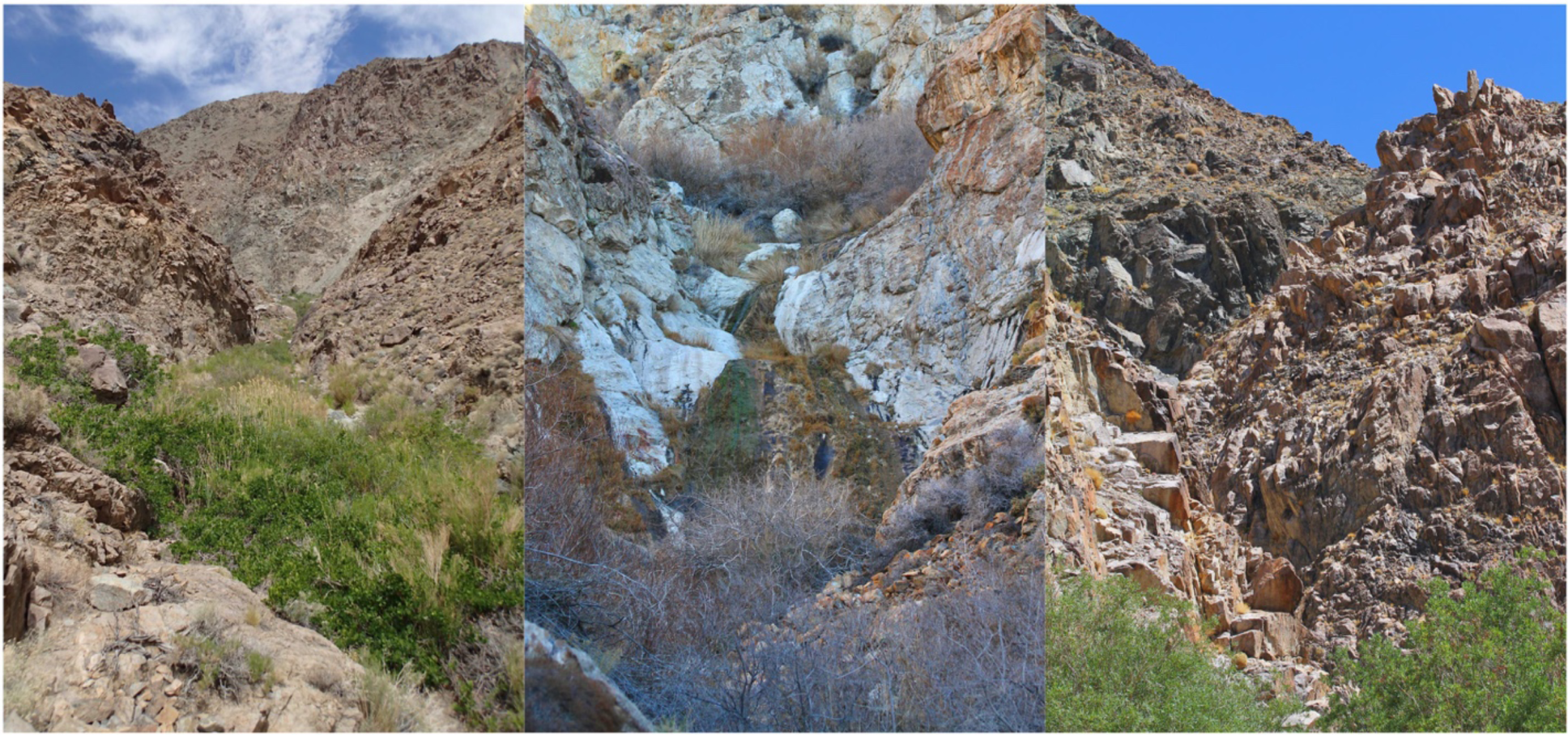
Habitat photos for *Elgaria panamintina* and/or *Batrachoseps campi* showing the rugged, rocky terrain and bedrock waterfalls that are common in occupied canyon-bottom microhabitat. Left to right: Water Canyon, Argus Mountains; unnamed canyon between Union Wash and Reward Mine, Inyo Mountains; French Spring, Inyo Mountains. Photographs by Adam G. Clause.

Here, we analyze the conservation status of *E. panamintina* and *B. campi*, using a dataset collated from white and gray literature, museum records, and contemporary field survey data. Our objective is to build a comprehensive threat analysis, generate a status assessment, and contrast our findings against outmoded sources and factual inaccuracies in the literature. We conclude by presenting management recommendations and research needs for both *E. panamintina* and *B. campi*, and highlight the necessity of ensuring scholarly standards in both technical and peer-reviewed literature.

## Methods

We reviewed the available literature on both *E. panamintina* and *B. campi* using their common and scientific names (and all synonyms) as search terms in the ISI Web of Science and Zoological Record databases. We also acquired copies of relevant uncirculated gray literature, in the form of reports prepared for resource management agencies. In addition, we included published sources such as species accounts from books (including field guides), the IUCN Red List, and NatureServe within our concept of relevant literature despite their often less scholarly nature. Although these reports and other works are usually held in lower regard than peer-reviewed publications, they contain a large proportion of available technical knowledge relevant to our study, and many were cited in the Adkins Giese et al. (2012) listing petition for both species. As such, we consider it imperative to consider these sources in our analysis.

Concurrently with our literature review, we queried the VertNet online portal to create a database of museum records, supplemented with data obtained directly from relevant museums (California Academy of Sciences, CAS; Museum of Vertebrate Zoology, MVZ; Florida Museum of Natural History, UF-Herpetology; and Natural History Museum of Los Angeles County, LACM). To georeference literature/specimen locality data, evaluate land ownership, and quantify the presence of roads, we used digital US Geological Survey 7.5-min 1:24,000 topographical maps published in 2012 and 2015, corroborated with the California Atlas & Gazetteer™ (DeLorme 2015).

We complemented this literature review with field survey data that we collected in 2000–2001 and 2009–2018. Our survey methodology primarily consisted of visual encounter surveys completed on foot, but we also road cruised occasionally. We surveyed each locality 1–10+ times, with a total survey effort of 3–100+ person-hours per locality. When on foot, we surveyed riparian vegetation and talus habitats in an attempt to detect surface-active *E. panamintina* or *B. campi*, often supplemented by flip-and-replacement of cover objects such as rocks and logs. During surveys we recorded all threats to either species, which we define as any anthropogenic or non-anthropogenic action or condition known or reasonably likely to negatively affect individuals or their habitat. Our definition of a locality corresponds to individual drainage basins or sub-basins, and every locality that we recognize is at least 1 airline km distant from the nearest portion of any other. At localities represented by point-source springs, we surveyed the length of the available habitat whenever possible. At localities represented by creeks or streams, impassable waterfalls or other barriers often prevented us from viewing habitat in upstream reaches. However, we consider our survey coverage of these long, linear localities sufficient to identify nearly all possible threats. Due to major access constraints higher in the remote, rugged reaches of many canyons, impacts from humans and their attendant infrastructure/animals are usually most intense near the canyon mouth (Figure 2). In keeping with these landscape-use patterns, we always covered the lower reaches of the creeks and canyons in our surveys. For all new localities and elevation records for *E. panamintina* and *B. campi* discovered during our surveys, we deposited vouchers at the LACM. These vouchers consisted of at least one of the following: whole-body specimen(s), genetic tissue sample(s), and digital photo(s).

After compiling this combined dataset, we first reviewed existing knowledge of the distribution and relevant natural history of each species, to provide appropriate context for evaluating threats. Next, we assessed all threats to these species that we identified during our field surveys or that were mentioned by Adkins Giese et al. (2012). We categorized these threats using the 5-factor analysis used by the USFWS for listing decisions. These are: (Factor A) the present or threatened destruction, modification, or curtailment of the species’ habitat or range; (Factor B) overutilization for commercial, recreational, scientific, or educational purposes; (Factor C) disease or predation; (Factor D) the inadequacy of existing regulatory mechanisms; and (Factor E) other natural or manmade factors affecting the species’ continued existence (ESA 1973). Because of the broad nature of Factors A and E, we further divided them into seven and two sub-factors, respectively. In total, we thus identified 12 discrete threats to one or both focal species.

For each of these threats, we ranked its severity on a scale from 0 to 3, with definitions as follows: 0 = not currently affecting known localities, 1 = currently affecting <20% of known localities, 2 = currently affecting 20–50% of known localities, 3 = currently affecting >50% of known localities. We consider the divisions of this ranking scale fine enough to be informative, yet coarse enough to be resilient to changes in threat rankings following the acquisition of new survey data.

## Results and Discussion

We surveyed 73% (24/33) of the documented localities for *E. panamintina*, and 81% (17/21) of the documented localities for *B. campi*. These were generally the most logistically accessible localities. We also surveyed 16 additional localities with appropriate riparian/talus habitat that is suspected, but not known, to support one or both species (Figure 4, Table S1). We did not detect either species’ presence at these additional sites, and instead only recorded the incidence of threats. Due to variable search effort and imperfect detection rates for these secretive species, additional surveys may show that they do occur at some of the 16 sites in which neither species was found. In all, we surveyed 63 localities (the numbers given above do not sum to 63 because *E. panamintina* and *B. campi* co-occur at 7 localities). We provide threat scores for *E. panamintina* and *B. campi* across all known localities in Table 1. Below, we describe our results for both species in three separate sections: geographic distribution, natural history, and threats. In each section we compare our results against claims made in the literature, to clarify discrepancies.

**Figure 4.**
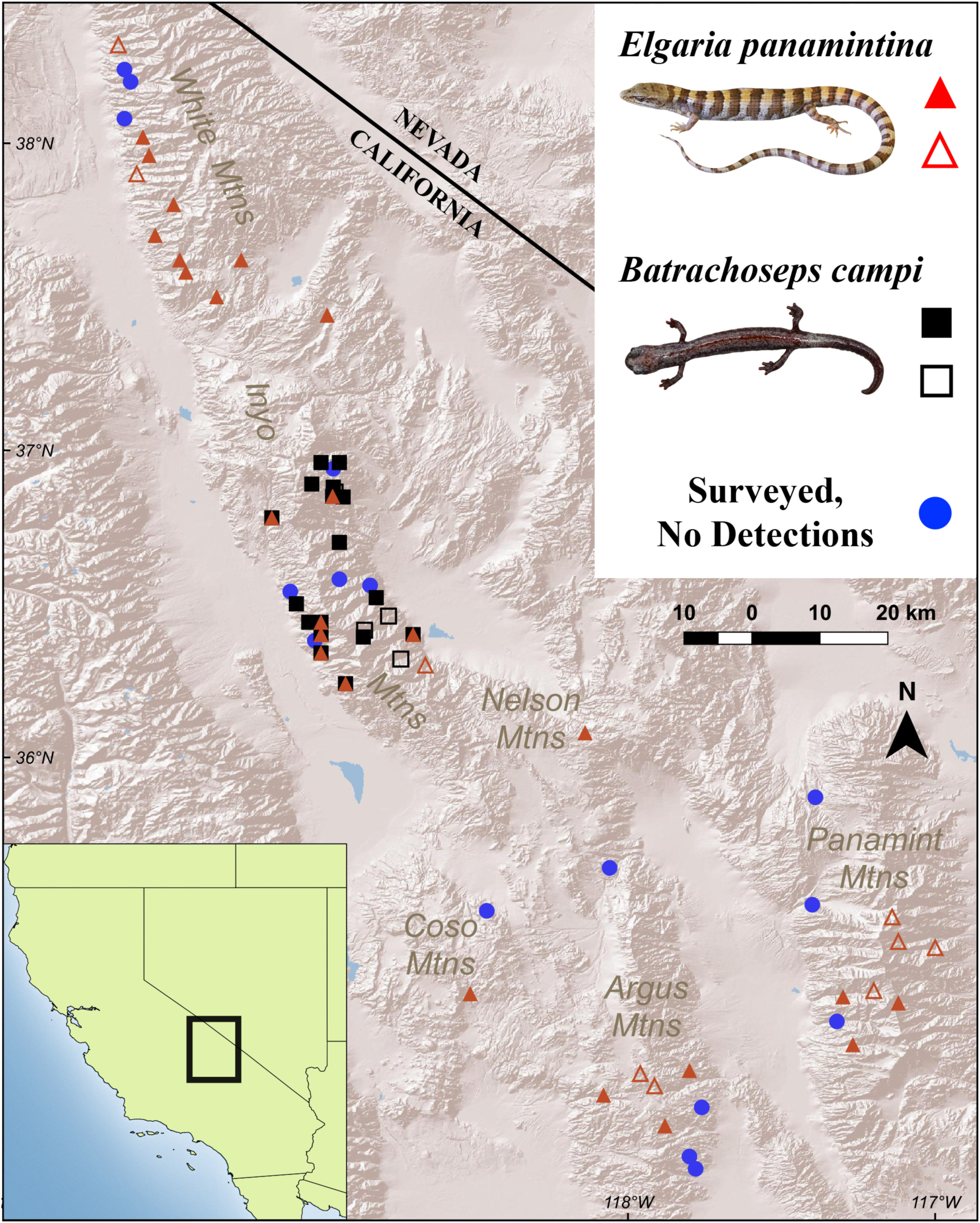
Rangewide locality-level distribution and survey coverage for *E. panamintina* and *B. campi*. Solid symbols show localities surveyed for this work, hollow symbols show historically surveyed localities.

**Table 1.** Rangewide threat scores for *Elgaria panamintina* and *Batrachoseps campi*. Score values correspond to the percentage of occupied localities currently known to be affected by the threat, as follows: 0 = 0%, 1 = less than 20%, 2 = 20–50%, 3 = over 50%. Scores marked with an asterisk (*) denote a prediction for the future; their current score is 0.

| Threat | USFWS<br>ESA Listing Factor | *Elgaria panamintina*<br>Score | *Batrachoseps campi*<br>Score |
| --- | --- | --- | --- |
| Water diversions | Factor A | 2 | 0 |
| Grazing by<br>feral/domestic<br>livestock | Factor A | 2 | 0 |
| Mining | Factor A | 0 | 1 |
| Roads and off-highway vehicles | Factor A | 2 | 0 |
| Invasive plants<br>(*Tamarix* spp.) | Factor A | 1 | 2 |
| Marijuana cultivation | Factor A | 1 | 1 |
| Renewable energy<br>development | Factor A | 0 | 0 |
| Overutilization | Factor B | 0 | 0 |
| Disease | Factor C | 0 | 0 |
| Inadequate<br>regulatory<br>mechanisms | Factor D | 0 | 0 |
| Climate change | Factor E | 3* | 3* |
| Flash floods | Factor E | 1 | 2 |

### Geographic Distribution

Across the western Great Basin and northern Mojave deserts of California, six named mountain ranges are known to support *E. panamintina*: the White, Inyo, Nelson, Coso, Argus, and Panamint mountains. In comparison, *B. campi* has a much more restricted distribution, and is known only from the Inyo Mountains (Figure 4). Banta (1965) predicted the occurrence of *E. panamintina* in three Nevada mountain ranges, Hammerson et al. (2005) included part of Nevada in their range map for the species, and Petersen et al. (2017) considered it “unconfirmed and potentially present” at two Nevada military installations. Nonetheless, no confirmed records exist for *E. panamintina* in Nevada, despite targeted survey effort in seemingly suitable habitat (J. Jones, Nevada Department of Wildlife, personal communication). Additionally, range maps by Hammerson (2004a, 2007) and Jockusch (2001) omit portions of the known distribution of *E. panamintina* and *B. campi*, respectively.

Elevation limits documented for *E. panamintina* range from 1,050 meters (m) (Surprise Canyon, Panamint Mountains; LACM PC 1738) to 2,330 m (Silver Canyon, White Mountains; LACM 187140). An imprecise record from Hunter Canyon, Inyo Mountains (LACM PC 2374) suggests that *E. panamintina* can occur below 600 m, but this remains unconfirmed. Elevation limits documented for *B. campi* are broader, ranging from 490 m (Hunter Canyon, MVZ 150363–66) to 2,625 m (Lead Canyon, LACM PC 2379). Although other authors present different limits for one or both species (Behler and King 1979, Jockusch 2001, Stebbins 2003, Mahrdt and Beaman 2009, Adkins Giese et al. 2012, Stebbins and McGinnis 2012), these sources either rely on outdated citations or do not present any supporting data or vouchers. Nevertheless, we predict that future surveys will expand the known elevation limits of both species.

A total of 33 localities are reported for *E. panamintina*, but nine are unvouchered and remain unverified (Figure 4, Table S1). Of these 24 vouchered localities, we report eight here for the first time. Across all 33 localities, ten are in the White Mountains (Dixon 1975; Stebbins 1985; Cunningham and Emmerich 2001), nine in the Inyo Mountains (Banta 1963; Giuliani 1977; Stebbins 1985; Macey and Papenfuss 1991b; Banta et al. 1996), one in the Nelson Mountains (Banta 1963), one in the Coso Mountains (Giuliani 1993), five in the Argus Mountains (Phillips Brandt Reddick Inc. 1983; Michael Brandman Associates Inc. 1988; LaBerteaux and Garlinger 1998; Morafka et al. 2001), and seven in the Panamint Mountains (Stebbins 1958; Anonymous 1982; Stebbins 1985; Banta et al. 1996; Cunningham and Emmerich 2001; Morafka et al. 2001). Thirty localities are from Inyo County, and the remaining three are in Mono County. For *B. campi*, a total of 21 localities are reported, but two remain unvouchered and unverified (Figure 4, Table S1). Of these 19 vouchered localities, we report two here for the first time. Fourteen localities are on the east slope Inyo Mountains, and the remaining seven are on the west slope (Giuliani 1977, 1988, 1990; Clause et al. 2014). All are from Inyo County.

Much confusion exists in the literature regarding the known locality-level distribution of both *E. panamintina* and *B. campi*. In the case of *E. panamintina*, some localities are considered independent by some authors, but actually are nested (e.g., Brewery Spring and Limekiln Spring lie within Surprise Canyon). Other localities are listed as separate, but actually represent a spatially proximate group of records best represented as a single locality (e.g., the many records from the southwestern CA Highway 168 corridor). Synonymous localities are also treated as different (e.g., Batchelder Spring = Toll House Spring), and some are incorrectly spelled (Westgard Pass misspelled as Westguard or West Guard), further adding to the confusion. These issues have led to repeated underreporting or overreporting of the true number of localities (Banta et al. 1996; Hammerson et al. 2005; Hammerson 2007; Mahrdt and Beaman 2009). Compounding problems are caused by authors overlooking gray literature or citing outdated sources, resulting in further underreporting of the distributions of one or both species (Jennings and Hayes 1994; Jockusch 2001; Hammerson 2004a, 2004b; Adkins Giese et al. 2012, Stebbins and McGinnis 2012). Mislabeled museum specimens have also contributed to one error for *B. campi* that we correct here. Specimens MVZ 150377–86, listed as originating from Pat Keyes Canyon and accepted by Clause et al. (2014) as the sole substantiation of that locality, were in fact collected at McElvoy Canyon as shown by a careful reading of Giuliani (1977) and his unpublished 1976 field notes. Our examination of Kay Yanev’s field notes for these specimens also demonstrate that McElvoy Canyon is the correct locality, and that Giuliani was the original collector.

All reported localities for both *E. panamintina* and *B. campi* lie entirely on federal land (Table S1). Rangewide, *E. panamintina* occurs on lands managed by the USDA Forest Service, Inyo National Forest (INF) (12 localities); U.S. Bureau of Land Management (BLM) (10 localities); National Park Service, Death Valley National Park (DVNP) (8 localities, of which 2 are shared with the BLM); and China Lake Naval Air Weapons Station (CLNAWS) (5 localities). Three of these federal agencies also manage lands that support all reported localities for *B. campi*: BLM (13 localities), INF (7 localities), and DVNP (1 locality). These lands are all under minimal or no development pressure, and they retain their natural character. We are unaware of any private land with habitat that is potentially suitable for *E. panamintina* or *B. campi*. This reality contradicts a statement by Jennings and Hayes (1994) that “all except two of the known populations of Panamint alligator lizard occur on private lands.” This statement was erroneous even in 1994, but has propagated widely across the literature (Hammerson 2007; Mahrdt and Beaman 2009; Adkins Giese et al. 2012; Stebbins and McGinnis 2012; Thomson et al. 2016). Furthermore, Adkins Giese et al. (2012) make an implicit claim, without supporting evidence, that at least one population of *B. campi* is on private land.

The known distribution of both *E. panamintina* and *B. campi* solely on public land directly relates to their population health and habitat quality, which have been misinterpreted in the literature. Jennings and Hayes (1994) state that populations of *E. panamintina* are experiencing “habitat loss,” Adkins Giese et al. (2012) indicate a “decline” in the species, and Hammerson (2007) states that there is a “probably continuing decline” in both population size and extent and quality of habitat. Similarly, Hammerson (2004a) indicates a “continuing decline” in number of mature individuals and in extent and quality of habitat for *B. campi*, Evelyn and Sweet (2012) claim that abundance of *B. campi* has “likely declined,” Papenfuss and Macey (1986) assert that spring diversions “likely” led to extirpation of “some populations,” and Adkins Giese et al. (2012) state that water diversion “causes extirpations” of this species. However, all of these claims are speculative and unsupported by data. Although it is possible that localized declines may have gone undetected, there is no evidence of population declines, population extirpation, or population-scale habitat conversion for *E. panamintina* anywhere in its range. For *B. campi* our survey work indicates that five localities experienced recent habitat loss and possible population declines due to flash floods, although the majority of *B. campi* localities that we surveyed have maintained their habitat and consequently, populations.

### Natural History

Habitat requirements for *E. panamintina* and *B. campi* are an oft-misunderstood aspect of their natural history. Both species are typically considered narrow habitat specialists found only in microhabitats immediately adjacent to perennial surface water (Stebbins 1958; Marlow et al. 1979; Papenfuss and Macey 1986; Jennings and Hayes 1994; Stebbins 2003; Hammerson et al. 2005). However, this narrative ignores or minimizes both early (Banta 1963; Giuliani 1977) and more recent (Morrison and Hall 1999; Cunningham and Emmerich 2001) data showing much broader ecological tolerances for both species, although far more data exist for *E. panamintina*.

To date, 12 independent observations exist for *E. panamintina* in arid rocky habitat 2.7–6.4 kilometers (km) from perennial surface water or riparian habitat. These observations are spread across seven localities, and ten are supported by a museum voucher (MVZ 75918, MVZ 150327–29, MVZ 227761, LACM PC 1835, LACM PC 1849, LACM TC 4376–77, UF-Herpetology 152976). The sightings include a mating pair (Morafka et al. 2001; LACM PC 1849) and a likely gravid adult female (LACM TC 4377), suggesting that at least some observations reflect the existence of breeding populations in these areas. The recognition that *E. panamintina* occupies habitats far from surface water is analogous to our understanding of habitat use in the closely-related central peninsular alligator lizard, *Elgaria velazquezi*. Endemic to the deserts of Baja California Sur, Mexico (Leavitt et al. 2017), *E. velazquezi* was hypothesized to occur only at isolated oases (Grismer 1988; Grismer and McGuire 1993) but is now known from multiple arid, rocky localities several kilometers from the nearest perennial surface water or riparian zone (Grismer and Hollingsworth 2001).

For *B. campi*, four specimens (MVZ 190989–92) were collected from an “antifreeze pitfall trap set beside [a] mossy opening in limestone” atop a ridge about 0.4 km from the nearest riparian habitat (Giuliani 1977). Although five additional anecdotal accounts exist for *B. campi* individuals claimed to be found in nearly identical microhabitats “far from water” in moist ridgetop crevices (Giuliani 1977), these anecdotes are unvouchered and unsubstantiated by biologists. We did not survey the vouchered locality for verification, although we did survey some sites with similar microhabitat characteristics elsewhere but did not detect *B. campi*. Despite the pitfall-trapped specimens demonstrating that *B. campi* can occupy habitat far from flowing water, occupancy rates in suitable, moist, non-riparian microhabitats remain unknown. We encourage additional survey effort in these non-traditional areas to better resolve this situation, while recognizing that these surveys will likely be challenging because such habitat is rarely found at the surface. Nonetheless, well-documented vouchered records do show that neither *E. panamintina* nor *B. campi* is restricted solely to areas with flowing perennial water or riparian vegetation, contrary to earlier stereotypes and recent misstatements by some authors (Adkins Giese et al. 2012; Stebbins and McGinnis 2012). Although we recognize that the presence of *E. panamintina*, and particularly *B. campi*, in these more arid habitats does not necessarily reflect the existence of self-sustaining populations with long-term viability, this uncertainty also applies to many mesic localities where these species occur. More data are needed on the metapopulation dynamics that might influence the suitability and population stability across these two habitat types for both species.

In addition to a lack of information on habitat preference, no rigorous, comprehensive estimates of population size or occupied habitat exist for *E. panamintina* or *B. campi* (but see Larson et al. [1984]). As such, these metrics for evaluating the species’ imperilment remain poorly quantified, and we instead use available data on the presence of suitable habitat, ongoing reproduction, and incidence of threats to infer population health elsewhere in this contribution. Estimates for *E. panamintina* by Hammerson et al. (2005) and Hammerson (2007), which are derived from an arbitrary assumption that 50 or more adults exist in each of 20 populations, yielded a total adult population estimate of at least 1,000 individuals. Hammerson et al. (2005) and Hammerson (2007) also provided a similarly coarse estimate of total occupied habitat of less than 5 km^2^ for *E. panamintina*, based on the arbitrary assumption of dimensions of 2 km X 0.1 km for each of about 20 occupied habitat patches. For *B. campi*, Larson et al. (1984) estimated a total effective population size of 14,000 across 12 populations, based on allele frequencies derived from allozyme data first published by Yanev and Wake (1981). Using field survey data from 12 occupied localities, Giuliani (1977) categorized *B. campi* habitat into four separate bins, reporting 11.9 linear mi of “excellent” and “good” habitat, and 10.3 linear mi of “poor” and “very poor” habitat. However, this analysis excluded substantial riparian habitat at those localities, due to access difficulties. Papenfuss and Macey (1986) subsequently produced estimates for 13 *B. campi* localities, 10 of which overlapped with those analyzed by Giuliani (1977). In their study, Papenfuss and Macey estimated 14.82 ha of “ideal habitat” for *B. campi*, but they did not specify their criteria for diagnosing such habitat. Adkins Giese et al. (2012) subsequently misinterpreted these studies and did not acknowledge their limitations, citing those works as support for their statement that occupied habitat for *B. campi* “totals less than 20 ha.”

### Threat Overview

Table 1 quantifies our assessment of 12 discrete threats of potential concern for *E. panamintina* and/or *B. campi*: water diversions, climate change, flash floods, grazing, roads and off-highway vehicles, invasive plants, illegal marijuana cultivation, mining, disease, renewable energy development, overutilization, and inadequate regulatory mechanisms. Below, we also offer a narrative account of these 12 threats, summarizing available data and contrasting our analysis with assertions in the literature. We particularly concentrate on Adkins Giese et al. (2012), because they present arguably the most liberal threat assessment for both species. We discuss the 12 threats in roughly decreasing order of severity, and we provide all raw threat occurrence data in Table S1.

#### Threat of Water Diversions

The hydrology of surface flows in the mountain ranges inhabited by *E. panamintina* and *B. campi* is not well studied. However, these flows appear to be driven by precipitation, which feeds groundwater cells that discharge as perennial springs above the regional groundwater level (Patchick 1964; Jones 1965; Bedinger and Harrill 2012). We are unaware of any evidence to suggest that ongoing regional groundwater pumping, such as the highly-regulated groundwater withdrawal in the Owens Valley floor (Elmore et al. 2003), is decreasing surface flow at any site occupied by *E. panamintina* or *B. campi*.

The threat of water diversion or other anthropogenic change to hydrology is generally mentioned only in passing in the literature associated with *E. panamintina* (Jennings and Hayes 1994; Hammerson et al. 2005; Adkins Giese et al. 2012; Thomson et al. 2016), and only one source gives a specific example of this threat affecting an occupied locality (Anonymous 1982). In contrast, Adkins Giese et al. (2012) feature this threat prominently in their discussion of *B. campi*, but cite Giuliani (1988) for support without recognizing the outdated nature of this source, similar to Hansen and Wake (2005). Giuliani (1988) described substantial degradation due to water diversions and grazing at Barrel Springs in the Inyo Mountains, a locality occupied by both *E. panamintina* and *B. campi*. However, our repeated surveys since 2013 indicate that the nearby mine is inactive, all water diversion infrastructure is defunct, grazing impacts are nonexistent, and the riparian zone has regenerated to at least half of the linear extent described by Giuliani (1988). We have observed multiple individuals of both *E. panamintina* (including a likely gravid female) and *B. campi* (including juveniles and a gravid female) at this locality on several visits since 2013, indicating the presence of reproducing populations (A. G. Clause and C. J. Norment, unpublished data).

Cumulatively, our surveys documented active diversion infrastructure at five localities in the White Mountains occupied by *E. panamintina*, and at five other localities possibly occupied by this species in the White, Argus, and Coso mountains (Table S1). Stretches of riparian vegetation up to 150 m in length were killed off at two of these sites, seemingly due to lack of sufficient moisture. Although we did not document active water diversions at any localities occupied by *B. campi*, we did observe defunct water diversion infrastructure at six *B. campi* localities (some syntopically occupied by *E. panamintina*).

Although available evidence indicates that active water diversions are not currently widespread among the known localities for either species, and have decreased in occurrence from historical levels in parallel with a decrease in regional mining activity (discussed subsequently), such diversions still pose a current threat to *E. panamintina* and a potential future threat to both species. We emphasize the need for ongoing management of this threat throughout the range of both *E. panamintina* and *B. campi*, particularly in the context of increased future water demand due to climate change (discussed next).

#### Threat of Climate Change

Recent climate models for California generally predict a hotter, wetter future climate statewide, but the direction and magnitude of these predicted changes fluctuates broadly depending on the model (Polade et al. 2017). According to one recent forecast, by 2060 the region inhabited by *E. panamintina* and *B. campi* will experience a mean temperature increase of ca. 2–3°C, and a mean precipitation increase of ca. 10–60 mm (Wright et al. 2016). However, future climate regimes in California could also bring more extreme droughts (Cook et al. 2015; MacDonald et al. 2016; Swain et al. 2018) and reduced summer monsoon precipitation (Pascale et al. 2017). Although both *E. panamintina* and *B. campi* survived a prolonged regional mid-Holocene drought (LaMarche Jr. 1973), any climate-related loss of precipitation-fed riparian habitat would almost certainly be a stressor on populations occupying those habitats. In addition, droughts would likely create pressure to initiate new agricultural and municipal water diversions from these springs and creeks, potentially exacerbating the loss of riparian vegetation. Adkins Giese et al. (2012) briefly mention climate change as a threat to both *E. panamintina* and *B. campi*, but do not discuss climate forecast variability. Few other authors mention climate change as a threat to either species, and most do so only in passing (Hammerson 2004b; Thomson et al. 2016).

In addition to large uncertainties surrounding California’s future climate, it remains unclear how severely the outcome of a hotter, wetter, yet more variable and thus drought-prone climate would affect *E. panamintina* and *B. campi* populations. Recent thermal modeling indicates that climate warming will likely depress the activity and energetics of arid-land lizards, but these studies predicted lower climate change-related extinction risk in anguids (which includes all alligator lizards) than most lizard families analyzed (Sinervo et al. 2010, 2017). In contrast, species-specific maximum entropy (Maxent) ecological niche models predict *E. panamintina* and *B. campi* to be at high and intermediate risk, respectively, of climate change creating conditions unsuitable for population persistence by 2050 (Wright et al. 2013). Nonetheless, it is unclear if riparian-zone populations would become extirpated or instead persist at lower population sizes in more restricted patches of non-riparian habitat. It is also unclear if populations inhabiting rocky areas far from riparian zones or standing surface water will become extirpated under those future climatic conditions, although again those conditions would likely be a strong stressor on those populations, particularly for *B. campi*.

Ultimately, we consider climate change to be perhaps the greatest potential threat to the long-term persistence of both species, both intrinsically and because it could worsen the stressors of water diversions (discussed previously) and flash floods (discussed next).

#### Threat of Flash Floods

Beaty (1963) suggested that flash floods occur regularly in the White Mountains, caused primarily by localized summer thunderstorms. Dramatic re-sculpturing of canyon topography and severe destruction to riparian vegetation are typical results. Across the ranges of *E. panamintina* and *B. campi*, forecasted wetter climate regimes in California could exacerbate the frequency and/or severity of these flash floods in the future (Modrick and Georgakakos 2015; Polade et al. 2017; Swain et al. 2018), but see Pascale et al. (2017) for alternative predictions. Hansen and Wake (2005) and Adkins Giese et al. (2012) discuss the threat of flash floods to *B. campi*, but only Cunningham (2010) mentions this threat for *E. panamintina*. Based on our surveys and several published sources (Giuliani 1990; Hansen and Wake 2005; Cunningham 2010), over the last 30 years at least seven thunderstorms have caused flash floods across eight Inyo Mountains localities occupied by *E. panamintina* and/or *B. campi*, plus an additional locality occupied by *E. panamintina* in the Panamint Mountains (Table S1). Additional flash floods, throughout the range of both species, likely remained undocumented during that period. However, based on our surveys and those of Giuliani (1996), following a documented flash flood *E. panamintina* has persisted at every locality and *B. campi* has persisted at most. There are three localities at which *B. campi* has not been detected in post-flood resurveys: the south fork of Union Wash (although *E. panamintina* has persisted there), Waucoba Canyon, and the middle fork of Willow Creek (C. J. Norment, unpublished data). Although recent flash floods have damaged or destroyed known habitat for *B. campi* and *E. panamintina*, our surveys also suggest that many known localities of both species are likely insulated from this threat because occupied habitat lies in side-canyon drainages too small to capture enough rainfall for scouring to occur. Furthermore, because heavy rainfall is a known behavioral cue for many organisms, including stream abandonment behavior in a few invertebrate taxa (Lytle 1999), *E. panamintina* and *B. campi* could possess behavioral mechanisms to help them escape flash floods. Regardless of possible mechanisms, ultimately flash floods represent a natural disturbance regime that both species have withstood for millennia. However, flooding could certainly act as a driver of local extinctions in a metapopulation dynamic, and we hypothesize that more frequent or extreme flash floods might exceed the recolonization or demographic capacity of both species to respond to this stressor in the future.

#### Threat of Grazing

Feral burros and feral horses have populated much of California’s desert wildlands for decades, primarily a legacy of abandoned stock associated with historic settlers and miners (Weaver 1974). The negative effects and widespread distribution of feral burros in DVNP were discussed by Sanchez (1974), and Giuliani (1977) subsequently documented extensive damage by feral burros to riparian zones at multiple *B. campi* localities in the adjacent east slope Inyo Mountains. Surveys at the *E. panamintina*-occupied Haiwee Spring in the Coso Mountains reported it as suffering “heavy” and “concentrated” use by feral burros (Woodward and McDonald 1979), and Giuliani (1993) later reported “over-grazing” by cattle and continued presence of feral burros at this locality. A review by Kauffman and Krueger (1984) demonstrated that intense grazing by non-native ungulates typically causes direct loss of riparian vegetation cover due to browsing, breaking, and trampling, accompanied by compaction and erosion of soils. Jones (1981) correlated these structural habitat changes with reduced lizard community abundance and diversity in Arizona. Reinsche (2008) subsequently reviewed additional studies that variously resolved both positive and negative effects of grazing on several lizard assemblages in arid and semi-arid landscapes. Although none of these studies involved alligator lizards or salamanders, we consider it reasonable that heavy grazing pressure is likely not beneficial to either *E. panamintina* or *B. campi* due to negative effects such as reduced vegetative cover, disturbance of microsites, and contamination of water sources.

Importantly, contemporary grazing severity at most localities for both species is reduced from historical levels, due to major removal efforts by federal land managers. From 1979–1981, the BLM removed over 1,500 feral burros from the east-slope Inyo Mountains (Papenfuss and Macey 1986). From the 1980s to 2005, the Navy removed 9,500 feral burros and 3,280 feral horses from CLNAWS lands. Navy removals are ongoing, to fulfill the CLNAWS Comprehensive Land Use Management Plan objectives of eliminating feral burros and maintaining a cumulative feral horse herd of 170 animals (U.S. Navy and Bureau of Land Management 2005). The BLM cooperates with the Navy in this effort, and has removed hundreds of additional feral ungulates from adjacent BLM lands known to support *E. panamintina*. Moreover, DVNP has engaged in control of feral ungulates since 1939 (Sanchez 1974). The Park Service has removed hundreds of burros from within DVNP; cooperatively implements burro control on adjacent BLM lands; has a long-term management goal of zero burros within the park; and plans to retire cattle from the Hunter Mountain allotment, which supports a known *E. panamintina* locality (National Park Service 2002).

The effects of these control efforts have been dramatic in many areas, although feral ungulates are far from being completely eradicated from the range of *E. panamintina* or *B. campi*. Our surveys documented grazing damage to riparian habitat at only two *E. panamintina* localities: one each in the Argus and Nelson mountains. Elsewhere in the Argus Mountains, on land managed by the CLNAWS and BLM, surveys by LaBerteaux and Garlinger (1998) indicated “low” or “moderate” feral burro grazing impacts at four additional *E. panamintina* localities that we did not survey. Nonetheless, anecdotal evidence suggests that grazing impacts could remain high in parts of the Argus, Nelson, Coso and Panamint mountains, which were comparatively under-represented in our recent surveys for *E. panamintina*. Ongoing removal efforts in these three ranges might be below annual recruitment rates, suggesting that populations of feral burros and perhaps feral horses could be on the rise in these areas (Tom Campbell, CLNAWS, personal communication). Moreover, funding constraints and deep-seated political controversy (e.g., Animal Welfare Institute [2012]) complicate the long-term management or eradication of feral ungulates (Crowley et al. 2017). Ultimately, current data on grazing severity are unavailable for many localities, particularly for *E. panamintina*. Nonetheless, our surveys found no evidence of feral or domestic ungulate grazing at any *B. campi* locality, and we consider it unlikely that this threat would cover a large portion of the species’ range due to the many inaccessible locations it occupies.

Adkins Giese et al. (2012) largely overlook data that indicate recent but variable reductions in non-native grazing animals on rangelands. Instead, they cite outdated secondary sources (Papenfuss and Macey 1986; Jennings and Hayes 1994) to support their claims that grazing is a major contemporary threat to both *E. panamintina* and *B. campi*. Adkins Giese et al. (2012) also incorrectly cite a third source (Mahrdt and Beaman 2002) by claiming that overgrazing “is” a threat to *E. panamintina* when Mahrdt and Beaman (2002) indicate only that it “could” be a threat.

#### Threat of Roads and Off-Highway Vehicles

Neither roads nor off-highway vehicles (OHV) are mentioned in the literature as a threat to *B. campi*, save for an unsubstantiated claim by Evelyn and Sweet (2012) that “road construction” is a likely contributor to declines. However, Adkins Giese et al. (2012) make several erroneous statements about the threat these factors pose to *E. panamintina*. They overlook contrary evidence to claim that OHV use “has increased significantly” in the Panamint Mountains, and that road “construction” threatens the species. For both claims, they cite only Mahrdt and Beaman (2002) for support, despite that source’s outdated nature and lack of supporting documentation. Adkins Giese et al. (2012) also mischaracterize a statement by Mahrdt and Beaman (2002), claiming that vehicular traffic “threatens lizard populations” when their source says only that it “could threaten lizard populations.”

Available evidence indicates that roads do pose an ongoing threat to *E. panamintina*, but no threat to *B. campi*. A two-lane paved road parallels or bisects occupied habitat at four known *E. panamintina* localities. At one of these localities, multiple road-killed *E. panamintina* have been documented (Morrison and Hall 1999; Cunningham and Emmerich 2001; specimens UF-Herpetology 152976 and LACM 189186). However, the sole patch of riparian vegetation (which the road bisects) and nearby roadside talus still consistently yield detections of this species 43 years after their discovery there (Dixon 1975). Furthermore, although data are limited, there is no indication of a decline in detection probability; our annual surveys of the spring-fed riparian habitat since 2013 have documented over two dozen individual lizards, about one-quarter of which were juveniles (A. G. Clause, unpublished data). Elsewhere in the range of *E. panamintina*, dirt access roads regularly approach the mouths of occupied canyons, but only at seven localities do dirt roads parallel and/or bisect riparian or talus habitat. Although grading and widening of three of these dirt roads in 2012 damaged riparian plants (Klingler 2015), our surveys indicate that much of the vegetation has since recovered. For *B. campi*, no paved road exists within 3 km of occupied habitat. Furthermore, only at four localities does a dirt road approach within 2 km of occupied habitat, and those roads never reach riparian zones inhabited by *B. campi*. At one locality (Barrel Springs), an old dirt road that closely approached occupied riparian habitat is now completely impassable to vehicles due to intentional placement of boulders in the roadcut.

The related threat of OHV use is even less consequential to *E. panamintina* and again a non-threat to *B. campi*. Many canyons where these species occur have multiple steep, bedrock waterfalls (Giuliani 1977) that restrict OHV passage (Figure 3). Except for one locality in the White Mountains (Redding Canyon), our surveys did not document evidence of unauthorized OHV use in or along riparian habitats occupied by *E. panamintina*. Contrary to statements made by Adkins-Giese et al. (2012), the severity of this threat has been much reduced from historical levels, due to targeted efforts by federal land managers. Over 15 years ago, the BLM prohibited all vehicular travel at the *E. panamintina* type locality in the Panamint Mountains (BLM 2001). Our surveys show that this canyon’s riparian zone has regenerated substantially in the absence of vehicular traffic, reclaiming much of the former dirt road that was a popular site for OHV enthusiasts. For *B. campi*, we are unaware of any OHV use at a known locality.

#### Threat of Invasive Plants

The only non-native plant mentioned in the literature, or that we identified during our surveys, as a threat to *E. panamintina* or *B. campi* is saltcedar or tamarisk, *Tamarix* spp. These shrubs or small trees can form dense monoculture stands in invaded riparian areas (Di Tomaso 1998); they have variable, but sometimes high, evapotranspiration rates that can potentially reduce surface water availability (Cleverly 2013; Nagler and Glenn 2013); and they are often correlated with elevated salinity levels in soil and groundwater, although causation has rarely been demonstrated (Ohrtman and Lair 2013). Research into the effect of *Tamarix* on lizard communities in the arid southwestern U.S. was reviewed by Bateman et al. (2013), and although no study involves alligator lizards, available research generally reveals a pattern of reduced lizard diversity and abundance in *Tamarix* stands relative to uninvaded riparian habitat. We are unaware of any studies exploring the effect of *Tamarix* on salamanders, but we infer that reduced surface water availability and elevated salinity levels would likely negatively affect *B. campi*.

LaBerteaux and Garlinger (1998), documented *Tamarix* at two known *E. panamintina* localities in the Argus Mountains, but noted that the plants were highly localized across the riparian habitat. DeDecker (1991) indicated a “widespread infestation” of *Tamarix* in low-elevation reaches of the west slope White and Inyo Mountains, where the plant had become a “serious threat to springs and seeps.” Adkins Giese et al. (2012) considered *Tamarix* a threat to *E. panamintina*, and cited DeDecker (1991) and Mahrdt and Beaman (2002) to support their position. However, Mahrdt and Beaman (2002) only paraphrase statements from DeDecker (1991) for support and present no novel data. In a subsequent work, Mahrdt and Beaman (2009) again mentioned invasive plants and *Tamarix* as a possible threat to *E. panamintina* without offering supporting evidence. In contrast, to our knowledge no published source identifies invasive plants or *Tamarix* as a possible threat to *B. campi*.

Although it was likely more abundant in the region historically as indicated by DeDecker (1991), our survey data indicate that *Tamarix* is currently neither a widespread nor severe threat to *E. panamintina* or *B. campi*, although it is a greater threat to the latter species. Our surveys documented *Tamarix* at ten localities in the Inyo Mountains, of which three were occupied by *E. panamintina* and seven occupied by *B. campi*. Of these ten localities, four support < 20 plants and appear to be in an early stage of colonization, three support established populations that were recently treated mechanically and chemically by the BLM with some success, and two support plants only at the canyon mouth far from habitat occupied by either species. Elsewhere within the range of *E. panamintina*, our surveys documented *Tamarix* at only one additional locality, where the plants were present low in the canyon far from occupied habitat. Cumulatively, there is little evidence that *Tamarix* or other invasive plants currently pose a substantial threat to *E. panamintina* or *B. campi*. This reality is attributable, in large part, to decades of *Tamarix* control efforts by multiple federal agencies. Nonetheless, without concerted management this threat could worsen in the near future given the capacity of *Tamarix* to colonize and spread.

#### Threat of Illegal Marijuana Cultivation

No literature source mentions marijuana grows as a threat to either *E. panamintina* or *B. campi*. However, since 2014 our surveys revealed three recently destroyed or abandoned marijuana grows in remote canyons: one at an *E. panamintina* locality in the east slope Argus Mountains, one at a *B. campi* locality in the east slope Inyo Mountains, and one in the Inyo Mountains at a locality that could support one or both species. At the grow site in the Argus Mountains, we observed chopping damage to mature willows, terracing of the slopes immediately adjacent to the riparian zone, compaction of leaf litter, and defunct water diversion driplines, with these impacts covering a 2-hectare area. Additional negative effects, such as other forms of streamflow diversion (Bauer et al. 2015) along with water and soil contamination from pesticide/herbicide application, are also probable. Installation of similar grows elsewhere is a future threat to the riparian habitat of *E. panamintina* and *B. campi*, particularly in isolated canyons otherwise exposed to minimal direct human activities, as has been found elsewhere in California wildlands (Butsic and Brenner 2016). Although we caution that clandestine activities such as illegal marijuana cultivation are inherently challenging to quantify, which complicates any assessment of their prevalence or severity, this threat warrants ongoing management attention.

#### Threat of Mining

Knopf (1912) described a widespread decline in mining activity across the Inyo and White mountains beginning in the late 19^th^ century. A review of Inyo Mountains mineral resources by McKee et al. (1985) indicated the general continuation of this pattern, albeit with periodic spikes in mining activity corresponding to rises in gold prices. Papenfuss and Macey (1986) subsequently reported 361 mining claims in the Inyo Mountains “filed in and around 13 canyons where [*B. campi*] is found,” some of which also support *E. panamintina*. However, these authors did not define the phrase “in and around,” nor did they indicate which mining claims were active, inactive, or not yet acted upon. Adkins Giese et al. (2012) list mining as a threat to both *E. panamintina* and *B. campi*, but they provide no examples to support their assertions nor do they acknowledge the limitations of the Papenfuss and Macey (1986) source, which they cite in their discussion of *B. campi*. Other authors (Hammerson 2004b, Hansen and Wake 2005) cite Papenfuss and Macey (1986) in a similar fashion, overlooking its outdated nature, particularly in the case of *B. campi*, because all known populations of the species now occur either within DVNP or the Inyo Mountains Wilderness, which was created in 1994. Although wilderness designation offers some protection for at-risk species and their habitats, valid mining claims existing prior to 1 January 1984 can legally be exploited; permitted activities include “where essential the use of mechanized ground…equipment” and “use of land for…waterlines” (Legal Information Institute undated).

In keeping with the general decline in mining noted by Knopf (1912) and McKee et al. (1985), our surveys suggest that mining has continued to decline across the known range of both species, and is not currently affecting the habitat of either. Although the large footprint of the active Briggs gold mine in the Panamint Mountains lies adjacent to riparian habitat that might be occupied by *E. panamintina*, our surveys revealed no active mines within 0.8 km of known *E. panamintina* or *B. campi* habitat—only abandoned ones. We regularly documented old mining-related debris among riparian habitat occupied by both species but we never observed water flowing from abandoned mines or mine tailings, suggesting minimal water pollution by mining-related contaminants. Nonetheless, legacy effects of mining in the region have not been well-studied. Despite the apparent regional decline in mining activity, it has not ceased completely and economic shifts in the supply/demand of gold, silver, and other minerals could increase regional mining pressures in the future. For instance, an application to re-open the Robbie Hoyt Memorial Mine at a known *E. panamintina* locality in the White Mountains, which proposes widening a dirt road that currently impinges on occupied riparian habitat, was recently submitted for review (Inyo National Forest 2017a). Furthermore, a controversial application for a large gold mine on the Conglomerate Mesa, Inyo Mountains, which was later formally withdrawn (Timberline Resources Corporation 2008), has also been recently re-opened (Silver Standard Resources Inc. 2016). Ongoing vigilance by land managers against potential future mining threats remains essential.

#### Threat of Disease

No authors claim that disease threatens *E. panamintina* or *B. campi*, and no documentation exists of wild individuals of either species showing outward signs of ill health. Nevertheless, the future and possible current threat posed to *B. campi* from the disease chytridiomycosis, which is caused by the fungi *Batrachochytrium salamandrivorans* (*Bsal*), and *B. dendrobatidis* (*Bd*), warrants consideration.

Due to its recent discovery (Martel et al. 2013), *Bsal* remains poorly studied. Although it has not yet been documented in North America, *Bsal* has devastated salamander populations in northern Europe (Yap et al. 2017). In the region where *B. campi* occurs, spatial models predict low to moderate habitat suitability for *Bsal* (Yap et al. 2015), and low to moderate salamander vulnerability to *Bsal* (Richgels et al. 2016), although the latter result could be an artifact of low salamander species diversity in the region. In comparison, *Bd* is better studied and has been correlated with enigmatic declines in terrestrial plethodontid salamanders (Cheng et al. 2011). Nonetheless, field and laboratory studies indicate highly variable *Bd* infection rates among this group of salamanders, to which *B. campi* belongs (Van Rooij et al. 2011; Moffitt et al. 2015; Mendoza-Almeralla et al. 2016). One study of *Batrachoseps attenuatus* revealed evidence of mixed susceptibility to *Bd* and no evidence of measurable declines in wild populations (Weinstein 2009), while a retrospective analysis of three species of *Batrachoseps* from insular populations revealed consistently low prevalence of *Bd* infection (Yap et al. 2016). Conversely, a retrospective study of *B. attenuatus* negatively correlated modern-day population persistence with time to first detection of *Bd* infection (Sette et al. 2015). The sister species to *B. campi* (*B. wrighti*; Jockusch et al., 2015) is known to be capable of infection based on a single *Bd*-positive specimen (Weinstein, 2009), but no other information relating to chytridiomycosis susceptibility exists for the *Plethopsis* subgenus of *Batrachoseps*, which includes *B. campi*. The deep evolutionary divergence of *Plethopsis* (ca. 40 MYA; Shen et al. 2016) coupled with the ecological extremes inhabited by its component species (Jockusch and Wake 2002) could limit accurate inference about the effects of chytridiomycosis on *B. campi* populations using data from non-*Plethopsis* congeners. Thus, *Plethopsis*-specific chytridiomycosis research is needed to evaluate the threat this disease might pose to *B. campi*.

#### Threat of Renewable Energy Development

Although the literature for *E. panamintina* and *B. campi* does not mention renewable energy development as a stressor, we consider it worthy of management attention for *E. panamintina*. Utility-scale solar projects have been proposed by the Los Angeles Department of Water and Power in the Owens Valley at the base of the Inyo Mountains, and although these proposals were later withdrawn or cancelled (Manzanar Committee 2015), if reopened in the future such projects could impact the lower edge of *E. panamintina* habitat on upper alluvial fans at canyon mouths. Furthermore, a major geothermal energy development project in the Coso Mountains is less than 10 km from a known *E. panamintina* locality, and potentially suitable talus habitat exists within the project footprint. However, this habitat has never been surveyed for the species, and the draft Environmental Impact Statement did not consider possible impacts to *E. panamintina* (BLM 2012). Despite a lack of evidence that *E. panamintina* is currently being affected by this or any other energy infrastructure, renewable energy is a growth industry in the California deserts (CA Senate Bill No. 2 2011; California Energy Commission 2014) and thus warrants ongoing attention as a potential future threat.

#### Threat of Overutilization

The issue of overutilization has received attention in the literature for *E. panamintina* and *B. campi*, but this attention has been speculative. For *E. panamintina*, Mahrdt and Beaman (2002, 2009) indicated that illegal collecting “may” threaten populations, and Adkins Giese et al. (2012) cited the former source as the sole support for their claim that illegal collection “likely threatens populations” of the species. Giuliani (1977, 1988, 1990) expressed concern about collector-driven disturbance to *B. campi* populations, but he did not specify the basis for those concerns.

To our knowledge, no major hobbyist market exists for any species of *Batrachoseps* or *Elgaria*. We are also unaware of any evidence that overutilization is a population-level threat to *E. panamintina* or *B. campi*. Furthermore, both species are likely inherently resistant to overutilization due to their secretive life histories, generally low detection rates (Giuliani 1977, 1996; C. J. Norment, unpublished data), and occupancy of remote, rugged habitats (Figure 1, Figure 2). Reported scientific whole-body collection of *E. panamintina* amounts to fewer than 50 specimens spread across 16 localities over a period of 60+ years. Similarly, reported whole-body collection of *B. campi* sums to fewer than 200 individuals across 17 localities over a period of 40+ years. Collecting at these levels over such extended time periods is unlikely to have an appreciable effect on population persistence (Dubois and Nemésio 2007; Krell and Wheeler 2014; Poe and Armijo 2014; Rocha et al. 2014; Hope et al. 2018). Furthermore, current lethal scientific collection of both species is strictly regulated, and generally allowed only when documenting a new locality (L. Patterson, CDFW, personal communication). Nonetheless, quantifying the magnitude of legal and illegal wildlife trade is challenging, and can be prone to underestimation (Salzberg 1996; Schlaepfer et al. 2005).

Our surveys documented no evidence of collector-driven habitat disturbance, and we encountered possible collecting equipment only twice: two plywood boards (one since removed by unknown person[s]) at an occupied *E. panamintina* locality, and wood roofing shingles at a locality occupied by both *E. panamintina* and *B. campi*. Importantly, none of these cover objects showed signs of recent disturbance during our repeated surveys, suggesting a lack of regular visitation. Furthermore, these two localities represent the most easily-accessible sites for these species. For this reason, these sites would potentially be those most strongly affected by illegal collecting pressure; yet, they support populations that repeatedly yielded captures of multiple individuals, including juveniles, during our surveys (A. G. Clause and C. J. Norment, unpublished data).

#### Threat of Inadequate Regulatory Mechanisms

Several sources mention the existing state and federal regulatory mechanisms that cover *E. panamintina* and *B. campi* in California, in generally positive terms (e.g., Hansen and Wake 2005; Thomson et al. 2016). However, Adkins Giese et al. (2012) downplay these government protections, characterizing them as “insufficient” and “inadequate.” Importantly, Adkins Giese et al. (2012) do not fully acknowledge the role played by these regulations in past management actions implemented specifically to mitigate several threats to *E. panamintina* and *B. campi*. As discussed previously, existing legal protections have directly motivated beneficial interventions on behalf of both species in recent decades, resulting in decreased levels of grazing, OHV use, and invasive *Tamarix* spp. in occupied habitat.

At the state level, under California Code of Regulations Title 14 Sections 5.05 and 5.60, there is a zero bag limit for *E. panamintina* and *B. campi* under sportfishing regulations, making it illegal to collect either species for recreational purposes. A recreational collecting moratorium also exists for all *Batrachoseps* salamanders in Inyo County. Moreover, it is illegal to commercially collect either *E. panamintina* or *B. campi* in California unless a biological supply house obtains specific authorization from the California Department of Fish and Wildlife (CDFW) to collect them for sale to bona fide scientific and educational facilities—an unlikely scenario for these species (L. Patterson, CDFW, personal communication). For over 20 years, *E. panamintina* and *B. campi* have also been administratively designated as Species of Special Concern by CDFW. This designation is intended to direct research and management toward enigmatic but likely imperiled species as a means to prevent more stringent future listing, but it does not directly regulate destruction of habitats or individuals (Jennings and Hayes 1994; Thomson et al. 2016). Also, the State of California has some jurisdiction over water diversions on federal land through the California Environmental Quality Act (CEQA). Under this statute, surface water cannot be legally diverted without a state permit; applications to divert are made though the California Water Resources Control Board, which would require a CEQA analysis (S. Parmenter, CDFW, personal communication). This process could provide an additional level of regulatory protection for *E. panamintina* and *B. campi*, as long as the State of California has the political will to prevent water diversions in sensitive habitat.

At the federal level, both species are currently designated as Sensitive by the BLM (BLM 2006), and Species of Conservation Concern by the USDA Forest Service (under FSM 2670). The BLM is mandated to manage Sensitive species and their habitat in a multi-use context, by minimizing threats affecting the species and improving habitat, where applicable (BLM 2008). The USDA Forest Service is mandated to develop and implement management objectives for Species of Conservation Concern and their habitat. This management is designed to ensure that the species maintain viable populations on Forest Service lands, and do not become threatened or endangered due to Forest Service actions (see FSM 2670). However, in their ongoing Forest Plan revision, the Inyo National Forest proposed to exclude *E. panamintina* from their Species of Conservation Concern list (Inyo National Forest 2017b), which would decrease management attention for over one-third of the species’ known populations.

Legal protection of *E. panamintina*, *B. campi*, and their habitats would be strengthened by ESA listing and would help compensate for possible lax enforcement or even the repeal of existing protections in the future. Nevertheless, historical regulatory protections have clearly improved the status of both species, and there is no signal to suggest that similarly beneficial management will cease in the near future.

## Conclusion and Recommendations

Although scientific information on *E. panamintina* and *B. campi* is limited, our literature review and threat analysis showsthat available information often contradicts or highlights uncertainty associated with historical and contemporary literature for both species. Although many knowledge gaps remain and additional data are needed, we consider the current conservation status of *E. panamintina* and *B. campi* to be relatively secure, although *B. campi* is comparatively more imperiled due to flash food impacts on habitat and populations. It is true that both species are California endemics known from comparatively few localities, which places them at increased risk for local stochastic extinction; however, all known populations occur exclusively on federally managed land that retains or is recovering its natural character under existing regulatory mechanisms and associated management efforts. Although it is possible that localized declines may have gone undetected, no evidence exists of population declines, population extirpation, or large-scale habitat conversion for *E. panamintina*, and habitat loss and possible population declines are documented at only five sites for *B. campi*, all due to flash floods.

Of the 12 threats to *E. panamintina* and/or *B. campi* that we identified in our review, only three appear to be currently important: water diversions, climate change, and flash floods. Available data indicate that water diversions actively threaten multiple populations of *E. panamintina* and formerly threatened some *B. campi* populations. Pressure to initiate new diversions might increase in the future under predicted climate change scenarios, or if federal regulations are relaxed regarding wilderness area protection or mining claim development. Shrinking riparian zones under a hotter, more drought-prone predicted climate is also a concern. Additionally, if climate change causes more severe and/or frequent summer thunderstorms, the resulting destructive flash floods could exceed the adaptive capacity of populations of both species to deal with this stressor, and flash floods seem to have caused recent declines at some *B. campi* localities. However, substantial uncertainty exists in the regional climate forecasts across the range of both species. Available evidence (albeit more substantial for *E. panamintina* than *B. campi*) suggests that populations can persist in more arid, rocky habitat far from canyon-bottom riparian zones, although such non-riparian habitat may be rarely occupied by *B. campi* and may support smaller, less stable populations compared to riparian habitat.

Based on available data, the nine remaining threats to both species (grazing, roads and OHV use, invasive plants, illegal marijuana cultivation, mining, disease, renewable energy development, overutilization, and insufficient regulatory protection) are neither widespread nor severe at this time. Threats due to mining have declined from historical levels, and threats due to grazing, OHV use, and invasive *Tamarix* spp. have been reduced from historical levels through targeted action by federal land managers. However, we emphasize that these nine threats could become more severe in the future, in part due to potential changes in the political and regulatory landscape. Ongoing stewardship by resource managers is necessary to appropriately safeguard populations of *E. panamintina* and *B. campi*.

Broadly, the results of our threat assessment are independently supported. Using a rigorous, transparent eight-metric risk assessment framework, Thomson et al. (2016) identified and scored 45 imperiled yet non-ESA-listed amphibian and reptile taxa across their entire known distribution in the United States (all in California). The cumulative threat scores for *E. panamintina* and *B. campi* ranked low (28^th^ and 19^th^, respectively) within this cohort of taxa, corroborating our assessment of the comparatively secure conservation status of these two species. Remarkably, of the 15 most threatened taxa identified by Thomson et al. (2016), only four were included in the Adkins Giese et al. (2012) ESA petition. This discrepancy alludes to what we perceive as a disconnect between scientific knowledge and the species selection process used by Adkins Giese et al. (2012). Although Thomson et al. (2016) was published four years after Adkins Giese et al. (2012), most of the same data and sources were available to and used by both sets of authors.

A similar, although less pronounced, disconnect also exists between current scientific knowledge and the IUCN Red List categorizations for *E. panamintina* and *B. campi*. The categorization for both species hinges on judgments that in some cases are weakly supported by contemporary data, an unsurprising result given that these species accounts were generated over a decade ago (Hammerson 2004a, 2007). For *E. panamintina*, the Vulnerable categorization rests on the assumption that there is continuing decline in the number of mature individuals, and that no subpopulation exceeds 1,000 mature individuals, fulfilling criterion C2a(i) (Hammerson 2007). The categorization of *B. campi* as Endangered rests on the assumption that there are continuing declines in habitat extent, habitat quality, and number of mature individuals, fulfilling criteria B1b(iii,v) and B2b(iii,v) (Hammerson 2004a). Appropriately, Hammerson (2004a, 2007) qualifies his judgments throughout, and makes it clear that they are inferences or projections. However, no rigorous, comprehensive estimates of historical or contemporary population size, or number of mature individuals, exist for either species. Nor is there clear evidence of widespread, critical declines in habitat extent or quality, particularly for *E. panamintina*. Furthermore, based on the available evidence, inferences or projections that widespread population declines have occurred are weak, for three reasons. First, all contemporary resurveys of historical localities have produced detections (with three exceptions for *B. campi*), a result that would be unexpected if widespread declines had occurred. Second, although available data are limited, there is no strong signal of decreasing detections at any locality for *E. panamintina*, although *B. campi* detections have declined at 5 of 14 localities for which we have resurvey data (including the three that lack detections during resurveys). Third, at localities with perhaps the most severe human impacts (Barrel Springs and CA Highway 168), repeated detection of juveniles indicates successful ongoing reproduction in these populations. As such, despite lacking comprehensive rangewide data, available information suggests that populations of both species are likely relatively stable overall, albeit with possible population declines in some *B. campi* populations for which we have resurvey data, due to recent flash floods. We thus re-evaluate both *E. panamintina* and *B. campi* as Near Threatened on the IUCN Red List (IUCN Red List Criteria Version 3.1), downlistings that we consider a positive development (see Mallon and Jackson [2017]).

Independent of their IUCN listings, we expect that the existing status of both *E. panamintina* and *B. campi* as CDFW Species of Special Concern and BLM Sensitive Species will motivate continued attention to their protection. Available evidence indicates that these status listings are warranted, and we suggest that they remain unchanged. We similarly advocate for the continued inclusion of *B. campi* on the INF’s Species of Conservation Concern list, and strongly recommend reconsideration of the proposed exclusion of *E. panamintina* from that list (Inyo National Forest 2017b). A recent analysis of the conservation status of *E. panamintina* on INF lands evaluated the species as being of “high concern” or “some concern” across all eight threat categories assessed (Evelyn and Sweet 2012). Our surveys indicate that the 12 known localities for *E. panamintina* on INF lands include some of the most at-risk populations rangewide. Half are affected by roads that bisect riparian or talus habitat, almost half are affected by ongoing water diversions, and two localities (Barrel Springs and CA Highway 168) are perhaps the most strongly human-altered of any known site. Furthermore, the localities that drain into the Owens and Chalfant valleys are among the most vulnerable to pressures from increased water diversions, due to the agricultural, ranching, and housing development in those valleys. For these reasons, we consider it imperative that *E. panamintina* be included on the INF’s Sensitive Species list, to promote continued management efforts by INF that will help forestall any need for more stringent listing status in the future.

All federal agencies that support populations of both *E. panamintina* and *B. campi* face challenges managing for these species in the context of multi-purpose land use and a changing political environment (Norment, in press). This reality can lead to unavoidable complexity and tradeoffs for many management actions, problems that are widely recognized among conservation practitioners (Hirsch et al. 2010; Roe and Walpole 2010; McShane et al. 2011). For instance, regional agricultural and municipal water needs are likely to increasingly conflict with those of *E. panamintina* and *B. campi*, control of feral horses and burros often contradicts deep-seated value systems held by some stakeholders, and mining-related economic development can clash with protection of sensitive riparian habitat. Furthermore, limited resources and competing goals may complicate implementation of management interventions, especially given the remote landscapes occupied by *E. panamintina* and *B. campi*. Scientific uncertainty, which we regularly identified in our threat assessment, will also necessitate adaptive management of these species (Runge 2011). While recognizing these challenges, we nonetheless offer four management recommendations designed to promote the long-term population viability of both species (Figure 5). Our recommendations are targeted toward preservation of sensitive riparian habitats that are critical not only to *E. panamintina* and *B. campi*, but also to co-occurring species of regulatory interest including the Inyo California towhee, *Melozone crissalis eremophilus* (federally Threatened, state Endangered) and desert bighorn sheep, *Ovis canadensis nelsoni* (BLM Sensitive and INF Species of Conservation Concern). We hope that this alignment with broader resource protection goals will increase the relevance of our recommendations in prioritizing future conservation action. First, we recommend that existing water withdrawals on federal lands be carefully enumerated and tracked, and that any proposal to initiate new water withdrawals be vetted using a detailed environmental impact assessment. Second, we recommend the continued reduction of feral burro and horse populations on federal lands, and the drawdown of permitted animal unit months in the Hunter Mountain cattle grazing allotment within DVNP. Third, we recommend continued control of *Tamarix* spp. on federal lands using appropriate control methods, with particular emphasis on localities where *Tamarix* eradication is feasible due to low plant abundance, while considering potential impacts to native species and habitat. Fourth, we recommend that any new proposal for mineral resource extraction on federal lands be vetted using a detailed environmental impact assessment, and that any mining-related destruction or degradation of riparian zones be carefully controlled. Attention to the other threats identified in our assessment is also important for conserving *E. panamintina*, *B. campi*, and their habitats. But we argue that focusing limited available resources on the management of water withdrawals, grazing, invasive *Tamarix*, and mining will likely maximize return on investment, and minimize the need for more strict regulation to protect these species.

**Figure 5.**
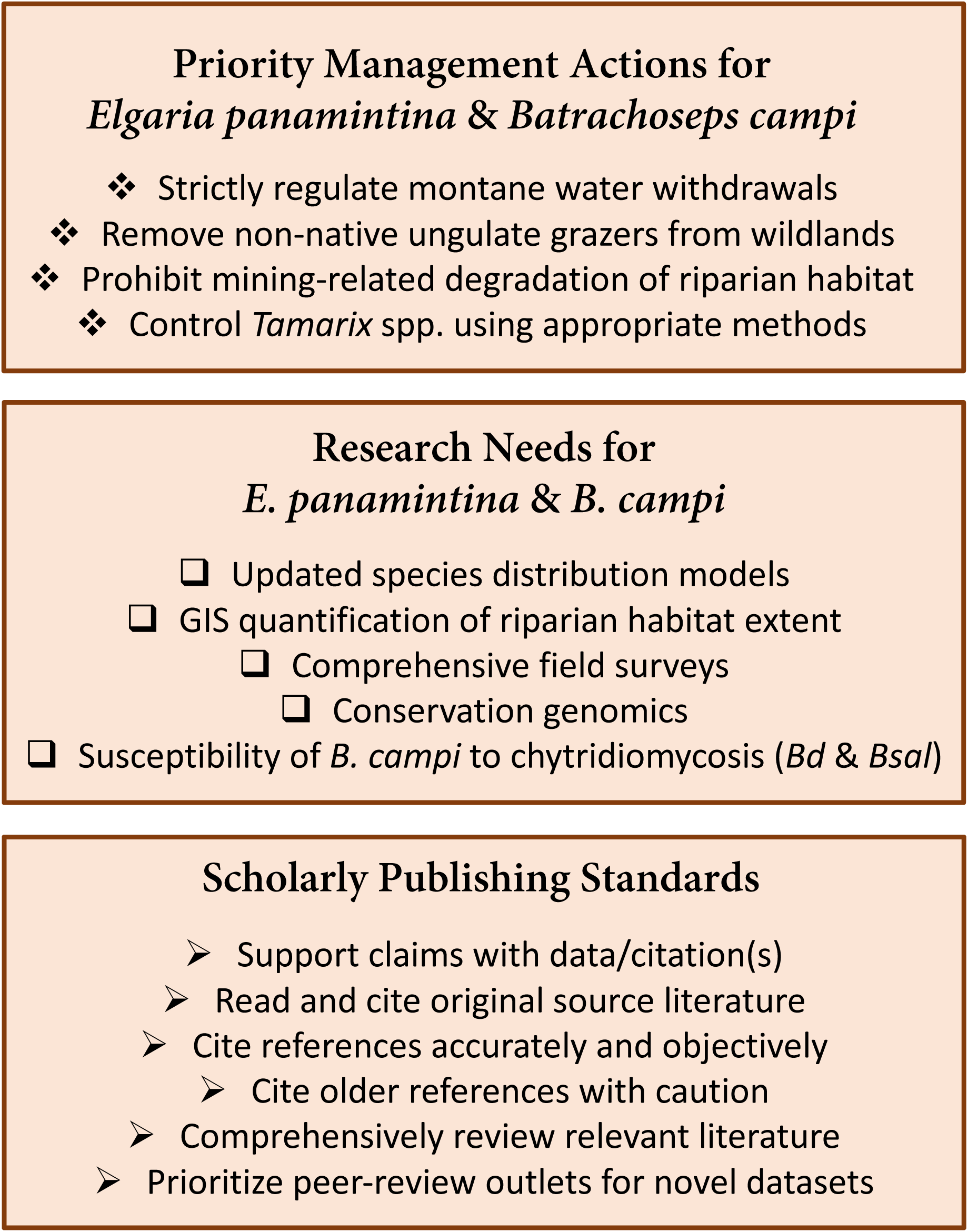
Recommendations for management and research relating to *Elgaria panamintina* and *Batrachoseps campi*, and reminder of scholarly scientific standards.

Through our status assessment and threat analysis, we also identified several research needs of immediate management relevance to both *E. panamintina* and *B. campi* (Figure 5). These include: (1) updated species distribution models, to inform targeted surveys of potential new localities and more rigorously evaluate the possibility of private-land populations; (2) rangewide GIS analysis of all riparian zones in the mountains occupied by these species, to produce a baseline assessment of habitat extent in the face of climate change; (3) comprehensive multi-day survey expeditions at suspected localities to verify and voucher species’ occupancy, ground-truth current threats, and evaluate habitat quality; (4) conservation genomics of known populations, to evaluate genetic diversity and estimate rates of gene flow among and between localities; and (5) field and/or laboratory studies of the susceptibility and prevalence of chytridiomycosis in *B. campi* populations from both *Bd* and *Bsal*. Some of these research needs have been advocated elsewhere (Thomson et al. 2016). Moreover, several goals (e.g., 3 and 4) are inherently linked and can be efficiently pursued simultaneously. We especially advise field workers in the Inyo Mountains to consider both *E. panamintina* and *B. campi* in their study aims, because these species likely co-occur at most riparian zones in that range. We encourage agency funding to support research on these topics, and emphasize that the results of such studies might lead to revisions of our threat assessment. Although our recent survey coverage included the majority of the known localities for both species, gaps do exist, particularly on lands managed by DVNP and CLNAWS.

Species-focused recommendations aside, our work additionally revealed the presence of several recurring scholarly problems that are of general interest to the broader scientific community, given the reliance of species status assessments on the best available science. The factual errors that we identified in the literature with respect to *E. panamintina* and *B. campi* are attributable to several causes, including: overlooking or selective use of available data, limited availability of some gray literature, the use of and failure to contextualize older data in light of more recent findings, misinterpretation or misrepresentation of data, and perpetuation of pre-existing literature errors. These problems are not unique, and have been identified elsewhere (Rubel and Arora 2008; Stromberg et al. 2009). In the *E. panamintina* and *B. campi* literature, the high frequency of errors could be a consequence of the relatively limited amount of data available in a peer-reviewed format, and limited accessibility of original data found only in uncirculated agency reports. For this reason, we encourage biologists and resource managers to prioritize the release of novel scientific data in a publicly accessible, peer-reviewed format whenever possible. Furthermore, we invite those who produce any scientific literature to strive for the following scholarly standards: (1) provide supporting evidence/citations for claims or statements; (2) reference original source literature, or scholarly review papers, when citing evidence for claims or statements; (3) cite references accurately, in a way that does not misrepresent the work of earlier authors; (4) cite older references with caution, and indicate when these might not reflect contemporary reality, and (5) comprehensively review the available literature on the topic of interest (Figure 3). Achieving these standards is a labor-intensive process, and no publication is ever perfect. Yet by striving to fulfill these scholarly guidelines (Figure 5), researchers will promote the best available science and help agencies tasked with resource protection to best prioritize their limited time and budgets. Furthermore, these recommendations will help to ensure that authors in any discipline will maximize the accuracy, value, and utility of their work, thereby assuring the integrity of the scientific community’s “bricks.”

## Acknowledgments

Author order follows the “sequence-determines-credit” approach as defined by Tscharntke et al. (2007). We thank B. Alexander, L. Bell, S. Brown, J. Calvin, T. Campbell, M. Dickes, J. Dittli, T. Hibbitts, S. Hillard, E. L. Jockusch, J. D. Hunt, C. Klingler, O. Klingler-Brown, J. Mattern, G. Milano, M. Murphy, D. Nielsen, M. T. Norment, S. P. Parker, S. Parmenter, K. Rademacher, D. Silverman, R. Smith, B. J. Thesing, D. York, and the late D. Giuliani for field assistance and data sharing. D. Pritchett at the University of California White Mountain Research Center, and C. Fidler at the University of California Berkeley Museum of Vertebrate Zoology Archives, graciously provided copies of the unpublished field notes of D. Giuliani and K. Yanev, respectively. We are grateful to N. Camacho and G. B. Pauly (LACM) for accessioning vouchers. D. J. House, M. R. Lambert, J. C. Maerz, and several members of the Maerz lab offered helpful comments on prior versions of this manuscript. D. LaBerteaux and C. Carter shared key literature. Financial support was provided through a University of Georgia Presidential Fellowship to AGC; a sabbatical leave granted to CJN by the College at Brockport, SUNY; and a contract awarded to CJN by the U.S. Fish and Wildlife Service. These funding sources had no role in influencing this research or its publication. Field work was authorized under California Department of Wildlife Scientific Collecting Permit #011663, Bureau of Land Management Permits #1110 (CA-170.31) P, #1110 (CA-6601.26) P, and #6500 (CA-170.31) P, USDA Inyo National Forest Permits #WMD15002 and #2300, USDI National Park Service Permit #DEVA-2017-SCI-0036, and University of Georgia IACUC AUP #A2012 10-004-Y1-A0, #A2016 02-001-Y1-A0, and #A2016 02-001-Y2-A0. A. Ellsworth issued a CDFW letter of authorization for survey work at the Indian Joe Springs Ecological Reserve. We dedicate this paper to the late David J. Morafka and Derham Giuliani.

## Supplementary Material

Table S1. Raw threat data and voucher information for all reported localities.

